# Prophage genomics reveals patterns in phage genome organization and replication

**DOI:** 10.1101/114819

**Authors:** Han Suh Kang, Katelyn McNair, Daniel A. Cuevas, Barbara A. Bailey, Anca M. Segall, Robert A. Edwards

**Affiliations:** Department of Biology, San Diego State University, 5500 Campanile Dr, San Diego, CA 92182; Department of Computer Science, San Diego State University, 5500 Campanile Dr, San Diego, CA 92182; Computational Sciences Research Center, San Diego State University, 5500 Campanile Dr, San Diego, CA 92182; Department of Mathematics and Statistics, San Diego State University, 5500 Campanile Dr, San Diego, CA 92182

## Abstract

Temperate phage genomes are highly variable mosaic collections of genes that infect a bacterial host, integrate into the host’s genome or replicate as low copy number plasmids, and are regulated to switch from the lysogenic to lytic cycles to generate new virions and escape their host. Genomes from most Bacterial phyla contain at least one or more prophages. We updated our PhiSpy algorithm to improve detection of prophages and to provide a web-based framework for PhiSpy. We have used this algorithm to identify 36,488 prophage regions from 11,941 bacterial genomes, including almost 600 prophages with no known homology to any proteins. Transfer RNA genes were abundant in the prophages, many of which alleviate the limits of translation efficiency due to host codon bias and presumably enable phages to surpass the normal capacity of the hosts’ translation machinery. We identified integrase genes in 15,765 prophages (43% of the prophages). The integrase was routinely located at either end of the integrated phage genome, and was used to orient and align prophage genomes to reveal their underlying organization. The conserved genome alignments of phages recapitulate early, middle, and late gene order in transcriptional control of phage genes, and demonstrate that gene order, presumably selected by transcription timing and/or coordination among functional modules has been stably conserved throughout phage evolution.

## Introduction

Phages are major drivers in global biogeochemical cycles through the control they exert on bacterial populations. Despite their abundance in the environment they are under-represented in sequence databases and knowledge-bases. Phages’ two life cycles, the lytic and lysogenic cycles, provide a unique window into phage genomics. During the lysogenic cycle, temperate phages integrate their genomes into their host’s genome in a non-lethal fashion to generate a region of DNA known as a prophage – either as a plasmid (1) or by directly recombining with the host’s genome (2, 3). The phage replicates in the host genome until external signals promote a transition to the lytic cycle, at which point the phage hijacks the host’s cellular DNA replication machinery in order to replicate the phage genome, and adopts the host’s protein synthesis machinery to create new phage particles. Eventually the phage lyse releases infective particles to find new hosts to attack.

The activation and movement of a prophage from one host to another enables phages to act as mediators of horizontal gene transfer, via either generalized or specialized transduction (4). This phage-mediated horizontal gene transfer may account for up to 10^15^ transduction events per second (5, 6). Phage-mediated horizontal gene transfer events are responsible for the spread of novel functions including the development of pathogenicity in bacterial strains such as *Escherichia coli* O157:H7 (5), *Vibrio cholerae* El Tor N16961 (7), *Streptococcus pyogenes* (8) and many *Staphylococcus aureus* strains (9). Phages may mediate the spread of antibiotic resistance in the mammalian intestine (10), and also transfer more benign genes such as those involved in photosynthesis (11).

We recently proposed a new model for phage host interactions that suggests at higher host densities ecological considerations favor the temperate lifestyle (12). The increased energy available at high host densities allows bacteria to support the replication of prophages, which in turn provide beneficial protection to the host via super-infection exclusion. The phages are using their host for replication, but are also providing protection for their host from other invading phages.

Identification of prophages in bacterial genomes is complicated by the problem that many phage genes do not share sequence homology with other proteins. In some cases a newly discovered phage does not carry any genes that share sequence similarity to any other gene in the available databases, highlighting how little we know about these viruses (13). The underrepresentation of well-annotated phage genes in available databases makes it difficult to assign functions to newly sequenced genomes. This is exacerbated as genome sequencing increases at a near-exponential rate, requiring reliance on automated annotation software that is dependent on primary sequence similarity to assign functions to new genes (14, 15). Most prophage identification algorithms rely on sequence similarity to known phage genes to identify prophage regions (16-20). This leads to a paradoxical problem of not being able to identify a prophage that does not contain any regions homologous to known phage proteins. Our prophage-finding program, PhiSpy, uses a novel algorithm that is capable of identifying a prophage even if there is no similarity to any of the proteins in the databases by analyzing a suite of genomic characteristics that separate phage-encoding regions from the bacterial core of the genome (21).

Here we present an analysis of over 11,000 bacterial genomes from which we identified 36,488 prophages. We used this data to identify a core phage genome organizational profile and demonstrate that a significant fraction of phage genomes adhere to this model. This genome structure allows us to predict the function of genes with no known homology solely based on their location in the genome. We identified the integrase genes in 15,162 prophages, and aligned those genes to infer phylogenetic relationships between prophages. Furthermore, we explored the connection between phage and host genomic characteristics to enable predictions of the bacteria that phages are infecting.

## Materials and Methods

### PhiSpy version 3.2 Improvements

Several improvements were made to the PhiSpy program, and we have released a new version of the software and a web site to provide an accessible interface to the program. The most important change we made was to allow PhiSpy to process genomes that were composed of multiple contigs, as the original version of PhiSpy would treat proteins from different contigs as adjoining proteins. The previous version of PhiSpy used a scanning window that looked 40 proteins ahead of the current protein being classified, but to remove the inherent direction bias, we changed the window to be 20 proteins on either side of the protein being classified. For proteins near the edges of contigs, which had fewer than 20 proteins on a side, we included all possible proteins. To update the list of prophage protein annotation keywords, all the phage genomes from PhAnToMe (http://www.phantome.org), and all the complete prokaryotic genomes from GenBank were downloaded. We compared the annotations of each set, and found 12 keywords unique to phage protein annotations: endolysin, terminase, baseplate, base plate, virion, antirepressor, excisionase, Cro-like repressor, cI-like repressor, rIIIA lysis, rI lysis, and rIIB lysis. These keywords were incorporated into PhiSpy for protein classification. In addition to these major changes, many other minor bug fixes were applied. A website was also built so that users can run PhiSpy online, without having to download and install the program and its dependencies: http://edwards.sdsu.edu/PhiSpy/. The source code for the updated version of PhiSpy is available from that website and from GitHub: https://github.com/linsalrob/PhiSpy.

### Dataset generation

All bacterial genomes were extracted from the SEED database, which contains both sequence and annotation information that is used by PhiSpy in making prophage designations (14, 15). The bacterial genomes contained a representative of every genus currently known. The updated version of PhiSpy (version 3.2) was run on all bacterial genomes using the generic bacterial training set option, and the outputs were parsed to identify all predicted prophage regions (Fig. 1). All PhiSpy predictions are available from https://edwards.sdsu.edu/PhiSpy and https://figshare.com/articles/PhiSpy_prophage_predictions/3146656.

**Fig. 1.**
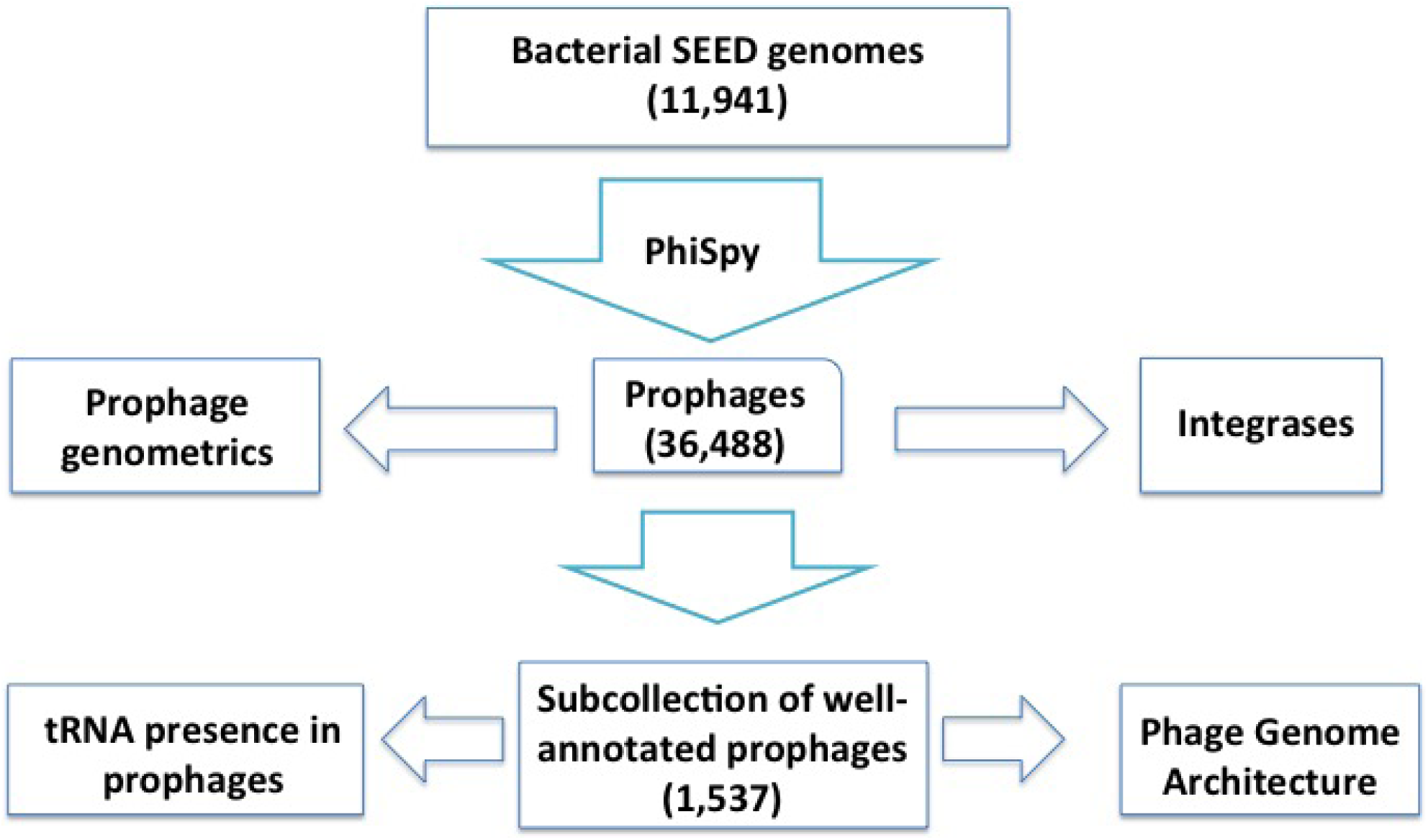
Workflow of the methodology used to identify prophages in complete microbial genomes.

### Raw sequence phage-host correlation studies

Prophage coordinates from PhiSpy were used to extract nucleotide sequence data from the original bacterial genome. The nucleotide sequences are available as FASTA files from https://edwards.sdsu.edu/PhiSpy and https://figshare.com/articles/PhiSpy_prophage_predictions/3146656. Putative tRNA-encoding genes were identified from those genomes using tRNAscan-SE (22). Modal codon usage was calculated from both the predicted prophage and the host genomes as described previously (23), and tRNA use and modal codon use were compared using the χ^2^-test. The G+C content of the phage and host nucleotide sequences were calculated using custom-written Python scripts.

### Genome heatmaps

High-quality prophage genome predictions were defined as those where at least one protein was annotated as an integrase, a capsid, a portal, a terminase, and a tail protein. The annotated genes in all the prophages were tabulated to identify a suite of genes that occur frequently in phage genome annotations (integrase, portal, tail, holin, transposase, terminase, endonuclease, capsid, antitermination, Holliday junction resolvase, baseplate assembly protein, primosomal protein, protease-like, lysin, toxin-encoding, and lysozyme).

To generate the genome alignment, each gene was assigned a unique letter indicating its function. A modified Clustal Omega alignment algorithm that scores +1 for matches and 0 for mismatches was used to align the annotated genomes.

### Integrase phylogenomics

A total of 24,936 integrase (Int) gene sequences were extracted from the high-quality manually curated dataset and clustered using CD-HIT (24) at a 90% 24,936 similarity threshold (an approximate virus “species” demarcation (16, 18) to produce 8,536 protein clusters. A representative of each cluster was used in the phylogenetic assessment of the 24,936 genes. In addition, short motifs were identified in the Int genes by using *k-mer* profiles for *k* from 4 to 7 amino acids long.

This approach allowed us to identify shared amino acid motifs including the eight-residue motif at the active site of the tyrosine recombinases. One hundred randomly selected integrases were aligned using Clustal Omega (25, 26) and visualized using FigTree (27). To compare the evolution of phage and bacterial proteins, all 24,936 phage Int proteins were aligned in a pair-wise fashion using the Smith-Waterman algorithm and the percent identity between each pair of proteins was calculated. Bacterial taxonomy was taken from the NCBI taxonomy database (28).

### Computational materials

Computational work was conducted on the Edwards Bioinformatics Lab compute cluster. Standard Python libraries were used for file parsing and Biopython version 1.65 was used to parse GenBank files. Scripts written for file parsing and data analysis are available in the github repository: https://github.com/hkang408/ProphageGenomics and from https://github.com/EdwardsLab.

## Results and Discussion

### Prophage genometrics

At the time of analysis, the SEED contained 11,941 bacterial genomes spanning 34 different phyla of Archaea and Bacteria (Fig. 2). A total of 36,465 prophages were identified in 9,883 bacterial hosts (82.76% of the bacteria have at least one prophage). Slightly fewer than one half of the prophages that were identified (15,765 prophages) contained an annotated Int protein, suggesting that they are phages with the potential to integrate. The mechanism of integration of the remaining phages could not be easily identified (see below). A total of 1,511,644 prophage ORFs were identified – of these, 45.7% or 798,235 ORFs had no known function. In addition, 567 genomic regions had no similarity to any of the proteins in the SEED non-redundant database (15) although these putative prophages had many other hallmarks of prophages such as shorter genes, many consecutive genes on the same DNA strand, and %GC compositional skews. A total of 7,341 portal genes, 6,228 holin genes, 14,771 capsid genes, and 59,010 tail-related genes were identified in the prophage dataset (Table 1).

**Fig. 2.**
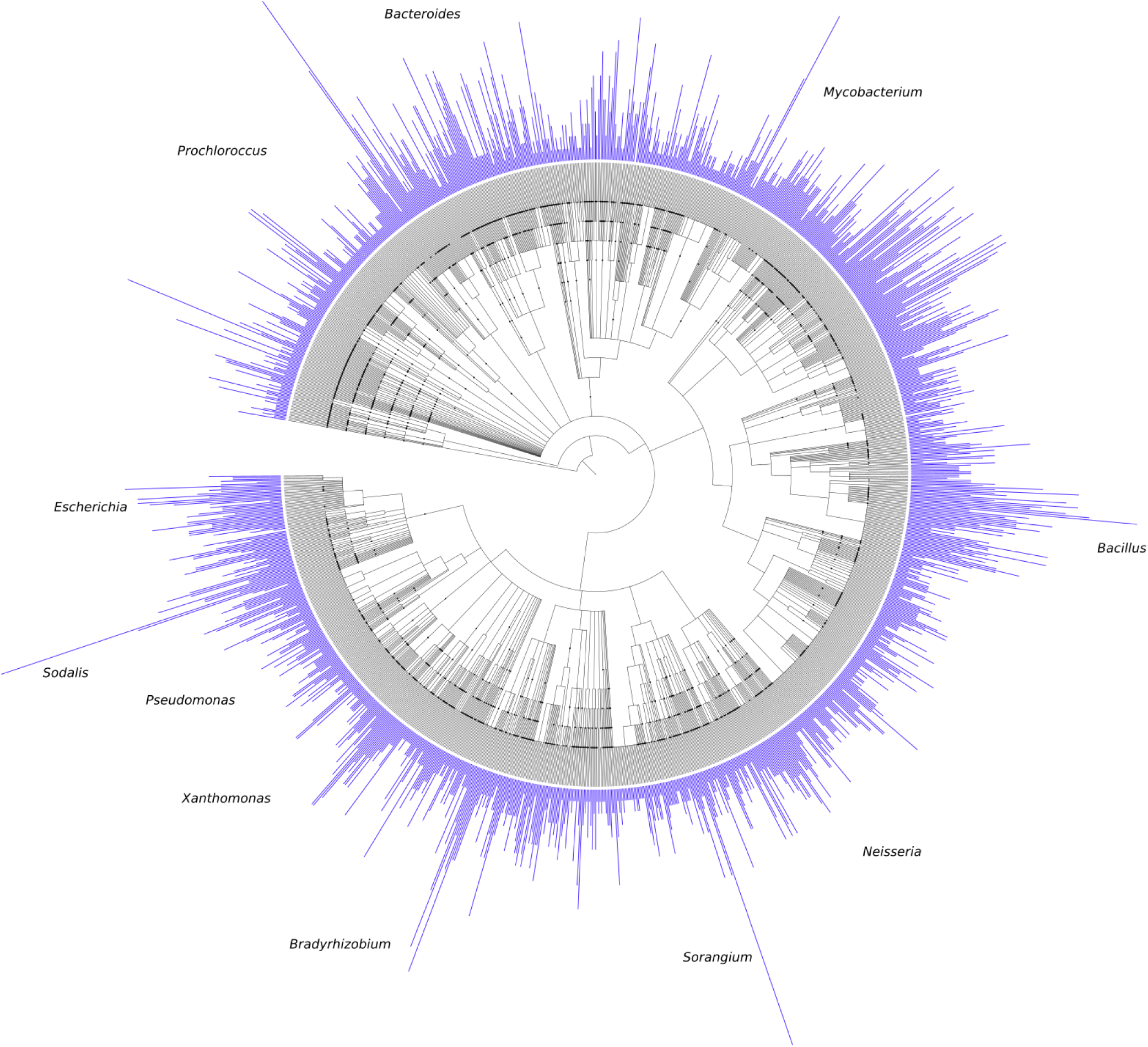
Bacterial genomes and the number of prophage regions per genome (blue bars). The phylogenetic tree is based on the NCBI taxonomy and was visualized using iTOL (http://itol.embl.de/). Selected genera have been annotated on the tree.

**Table 1.** Frequency of identification of different genes among the prophages. ORFs were identified from 36,465 prophages.

| **Function** | **Number of observations** |
| --- | --- |
| Integrase | 24,936 |
| Single-stranded binding protein | 2,556 |
| Endonuclease | 4,043 |
| DNA Polymerase | 5,011 |
| Protease | 6,575 |
| Terminase | 15,657 |
| Portal | 7,341 |
| Capsid | 14,771 |
| Tail-related | 59,010 |
| Baseplate | 3,728 |
| Holin | 6,228 |
| Lysin | 5,983 |
| Toxic | 132 |
| Transposase | 5,139 |
| Hypothetical | 737,992 |
| Other | 645,956 |
| **Total** | **1,511,644** |

As we have shown previously, some families of genes are easier to identify than others, and gaps remain in our understanding of families of core phage proteins that we cannot readily identify from sequence homology alone (29–31).

Across the most highly represented phyla, there were an average of between 2.7 (Actinobacteria) and 3.8 (Proteobacteria) prophages found per genome. Within phyla there were also differences (although not statistically significant) in the prophage genome content. For example, *Sodalis glossinidius,* a proteobacterium, had the most prophages at 29, while several phyla (Caldiserica, Deinococcus–Thermus, Dictyoglomi, Elusimicrobia, and Fibrobacteres) had no prophages, Spirochetes had the shortest prophage genomes on average (25,595 bp), and Bacteroidetes had the longest (46,059 bp) (Fig. 3). Across all of the prophages, the mean and median prophage genome length was 36,804 bp and 32,352 bp, respectively. Shorter prophage genomes may indicate that the phage is no longer viable and is in the process of being degraded or removed from the host’s genome (32). Alternatively, different genome lengths may represent different packaging strategies used by prophages associated with different phyla (33).

**Fig. 3.**
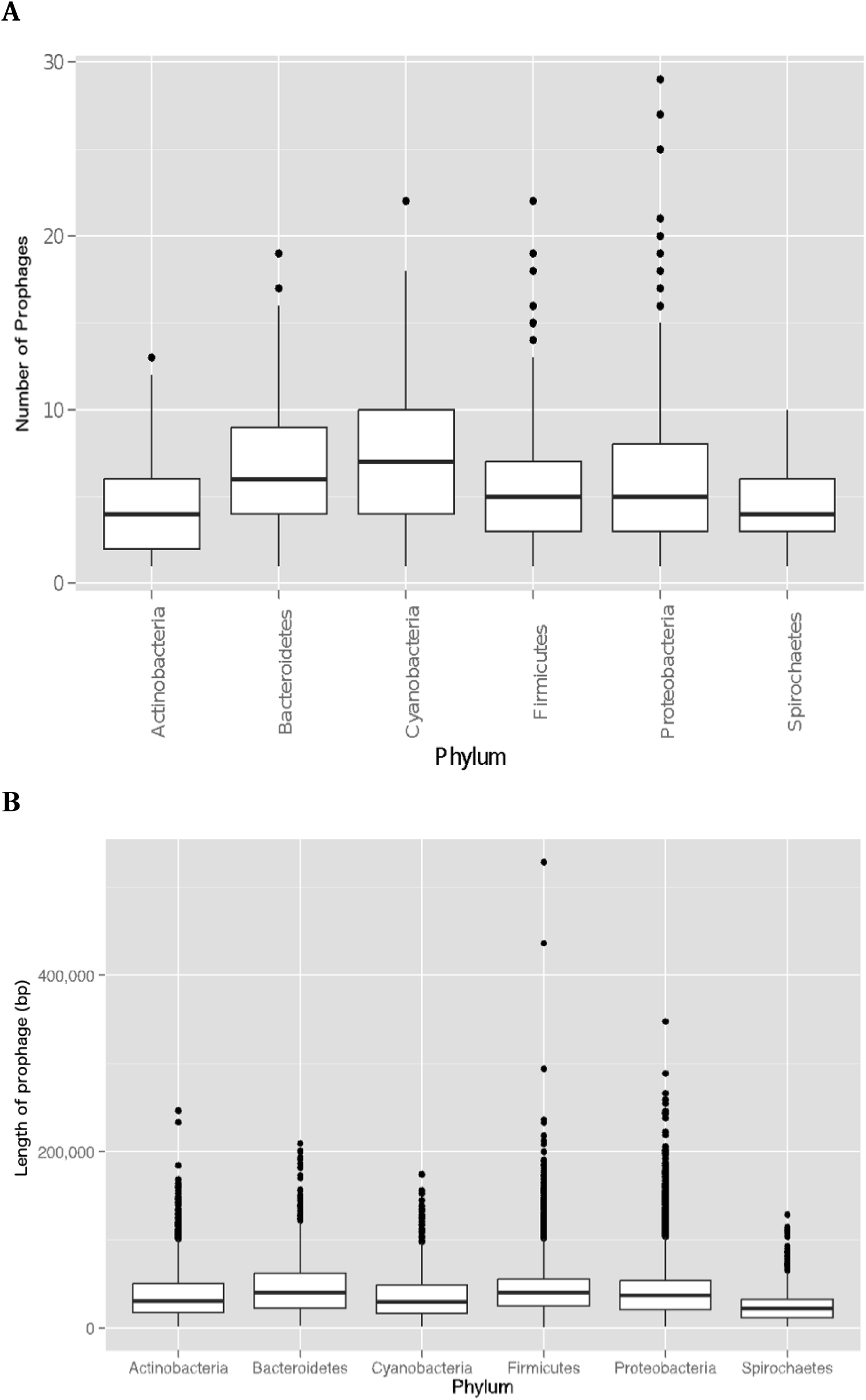
**(A)** Box and whisker plot denoting number of prophages identified per host, grouped by six phyla with most prophages. **(B)** Length (bp) of prophages identified grouped by six phyla with most prophages. In both A and B plots, the center line denotes the median of the distribution while box edges denote one standard deviation above or below the median.

Several prophage genome characteristics correlate with the host's characteristics (34). For example, the prophage’s and host's GC content correlated strongly (R^2^ = 0.95; Fig. 4). As we have shown previously, the GC content of phage genomes can be accurately used to predict the phyla of the host associated with a phage (to >95% accuracy), however GC content can not be accurately used to predict the phages’ host at lower taxonomic levels (35). Even though %GC content correlates at the phyla level, there are not enough degrees of freedom in %GC calculations to separate thousands of phages and their hosts. Percent GC is basically a *k*-mer DNA profile with *k* = 1. Higher *k*-mer DNA profiles increase the correlation between phage and host, but when *k* exceeds 7 nucleotides the data become sparse and over fit the predictions (35).

**Fig. 4.**
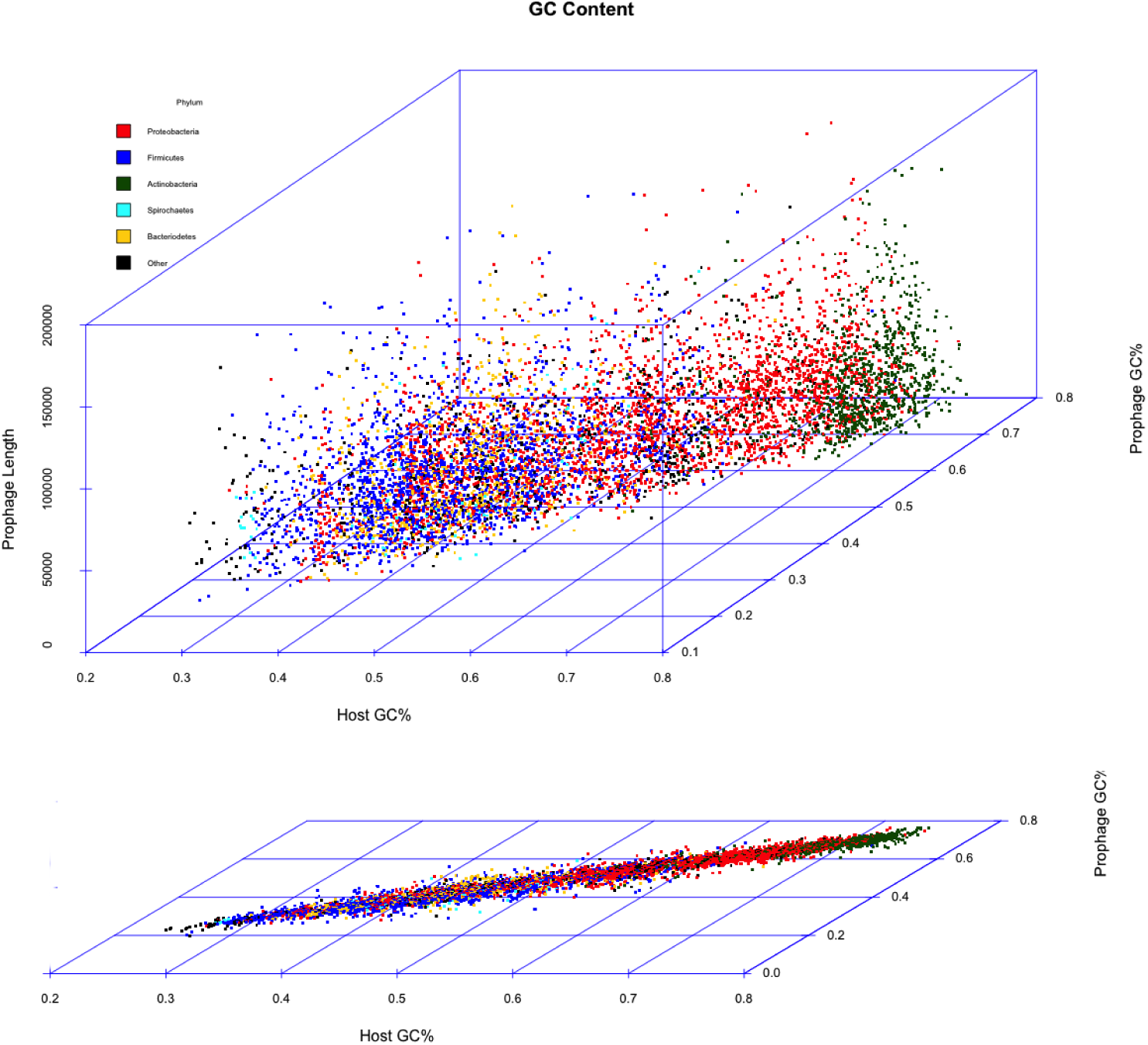
The %GC content of prophages shows strong correlation to %GC content of their hosts (R^2^ = 0.94735). There is no correlation between prophage %GC content and prophage length (upper panel).

### Phage integrase

Phage integrases are responsible for catalyzing the site-specific recombination event that inserts the phages into the hosts’ genome (36–41), as well as during replication to help in phage and plasmid genome maintenance. A total of 18,352 prophages (43%) contained an integrase-like protein from one of the two main families – the tyrosine and serine recombinases – named after the catalytic residue involved in recombination. The tyrosine recombinases include the well-studied integrase employed by phage λ to insert its own genome into the bacterial chromosome (2), while the serine recombinases are exemplified by the integrases of *Streptomyces* phage phiC31 (42) and *Mycobacteria* phage Bxb1 (43). The 24,936 integrase sequences were clustered at a 90% sequence identity threshold using CD-HIT, which resulted in 8,536 clusters. Representatives from each cluster were used to generate a smaller dataset with which to work.

Five-mer profiles of the amino acid sequences were used to identify an eight-residue motif in one of the catalytic region of tyrosine recombinases with a consensus sequence of **HDLRHTFA** (Fig. 5A). The very strong conservation of five of the amino acids (bolded) suggests that they participate in the catalytic activity of these recombinases. Several of these amino acids were identified earlier as invariant among tyrosine integrases or nearly so (44–46), and both of the histidines, the leucine, and the arginine have been experimentally demonstrated to participate in DNA recognition, binding, and/or catalysis (41). Alignment of one hundred randomly selected tyrosine integrases from the condensed integrase set shows that the proteins that contained the motif were grouped together (Fig. 5B), offering a potential route to identify tyrosine recombinases.

**Fig. 5.**
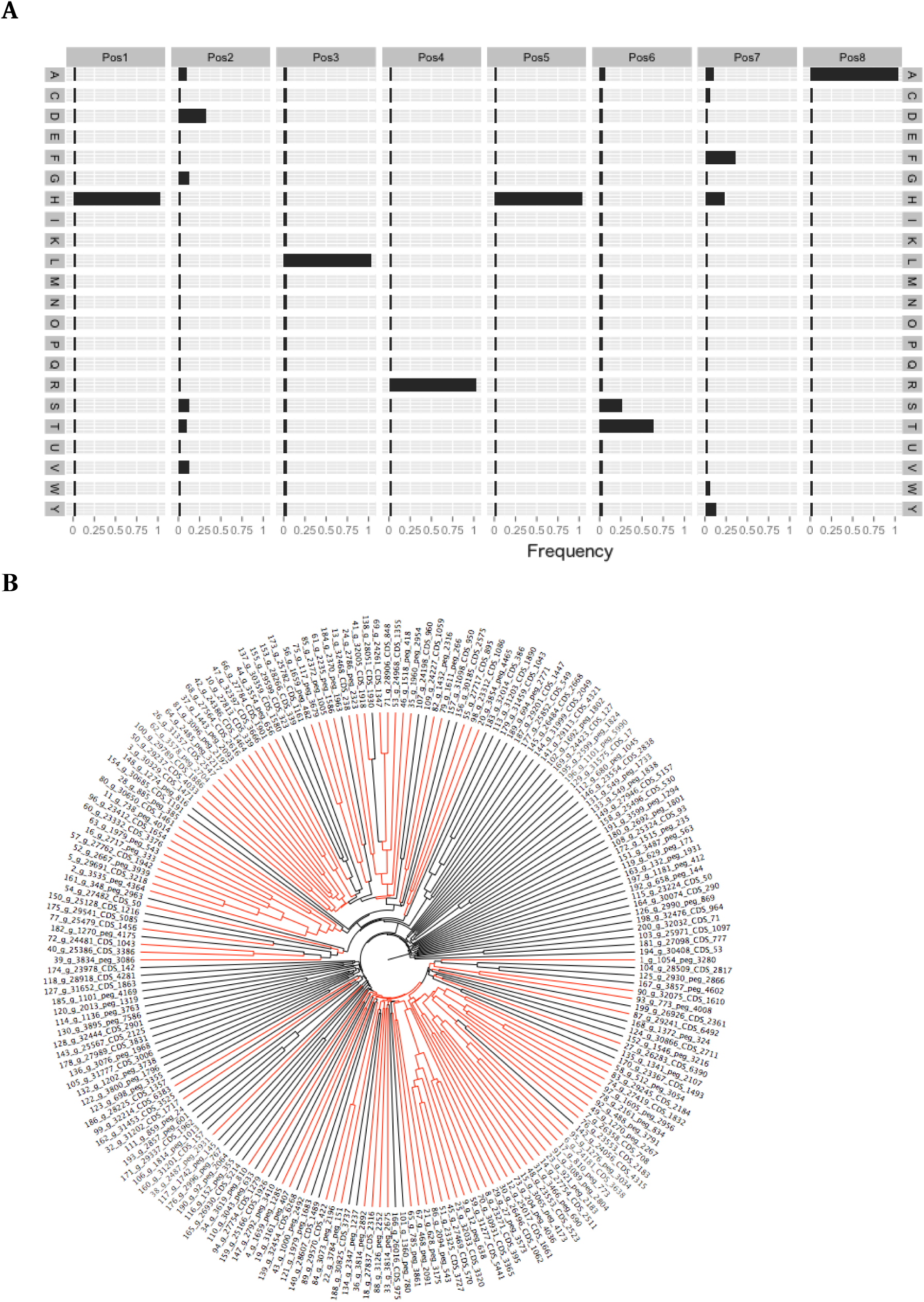
**(A)** Residue frequency of the eight-mer motif contained in the catalytic domain of phage HP1-like integrases. The bars represent the frequency with which each amino acid was found at each of the eight positions in the motif. The consensus sequence is HDLRHTFA. **(B)** A phylogenetic tree built from one hundred randomly selected integrases shows clusterings of integrase containing the eight-mer motif (highlighted in red).

### tRNA presence in prophages compared to tRNA integration occurrences

Many phages encode tRNA genes, which enhance phage fecundity (47, 48). Several hypotheses have been proposed to explain the presence of tRNA genes in phage genomes, including providing tRNAs to compensate for missing or weakly expressed genes in the host (49). A previous study of 37 phage genomes from 15 bacterial genomes demonstrated that phage tRNA genes compensate for codon bias between host and phage, especially when a codon is common in the phage and rare in the host (50). In addition, tRNA genes carried by the phage may replace those interrupted during phage insertion into the genome at tRNA genes (51, 52). It is also possible that encoding multiple tRNA genes provides the phage genome alternative attachment sites to integrate into the host's genome in case the preferred attachment site is mutated and/or occupied by another phage.

A set of 17,539 tRNA genes were identified from the phages in our total dataset and condensed to a subset of 3,833 tRNA genes (as described for the integrase genes). As expected, the tRNA gene profiles in the phages reflect the number of synonymous codons for each tRNA (Fig. 6). However, we found that tRNA-Met is one of the most commonly carried tRNA genes among the prophages we studied. Phages often encode a peptide deformylase enzyme that removes the formyl group from newly synthesized proteins (53, 54). We propose that the methionine tRNA genes encoded by the prophages are included to increase the overall rate of translation when the phage enters the lytic cycle. Together, the peptide deformylase and tRNA-Met suggest that phage protein synthesis saturates the host's protein synthesis machinery and that, in many cases, optimal production of phage progeny requires protein translation capacity which exceeds that provided by the host.

**Fig. 6.**
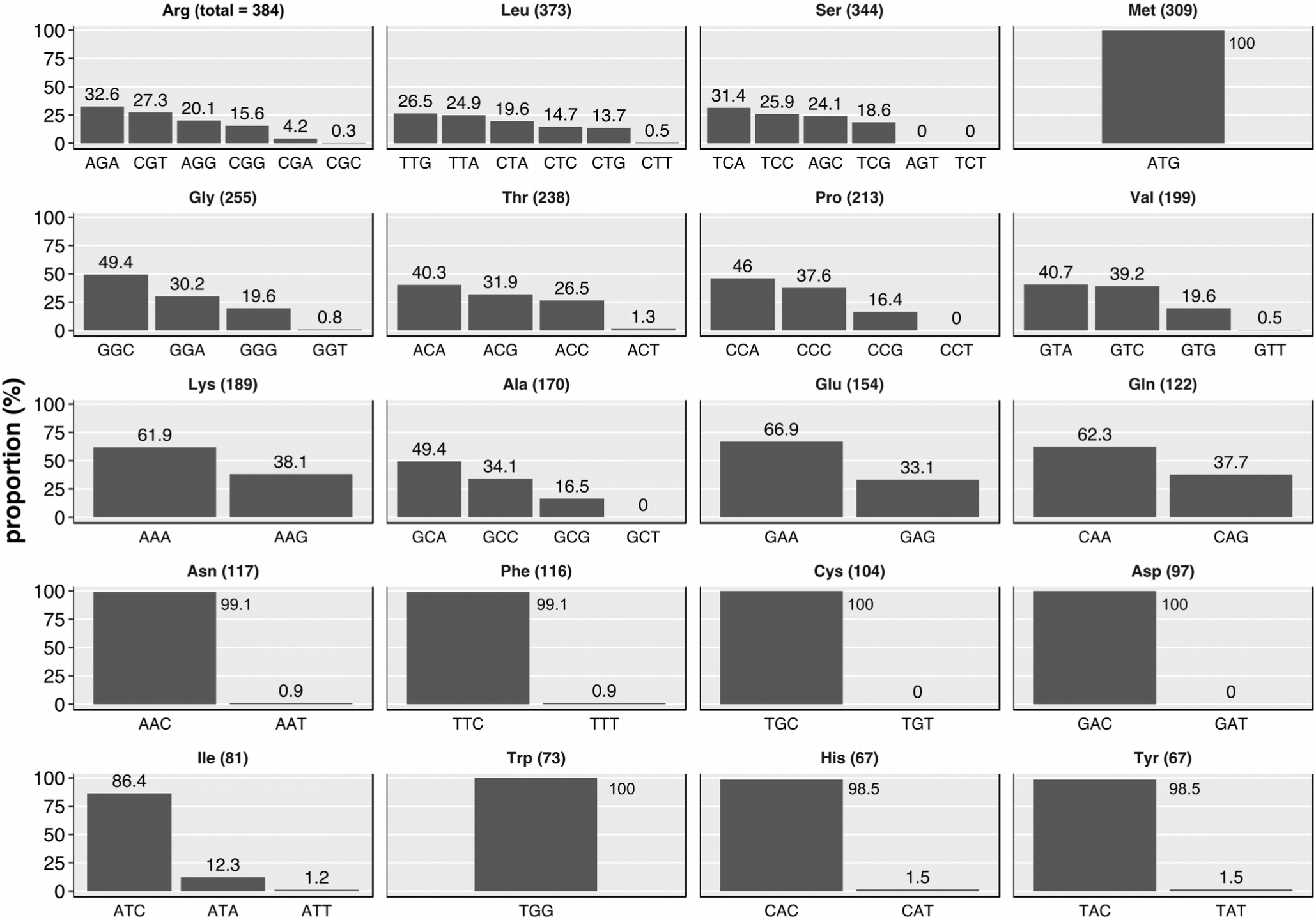
The frequency of different tRNAs identified within prophage regions. For each amino acid, the total number of phage-encoded tRNA is shown in parentheses above each graph. The percentage of those tRNAs for each of the possible anticodons is shown in the figure.

We explored the hypothesis that the tRNA genes compensate for codon bias between the host and the phage. For a tRNA gene found in a prophage, the anticodon was converted to the genomic DNA codon it’s associated with. This codon’s modal usage was calculated for the prophage’s genome, as well as for the host’s genome without the prophage sequence. The codon’s modal usage in the phage and host was compared using the χ^2^-test. We found that 41.3% of the codons associated with the tRNA found in a prophage had statistically higher modal usage in the phage than in the host (Fig. 6).

We applied the same approach to compare the phage’s modal usage of the same codon with an expected usage assuming an even distribution given the number of synonymous codons available. This comparison revealed that 31% of the codon associated with the tRNA found in a prophage had a higher usage in the phage than expected by chance. Together, these results suggest that phage-encoded tRNA are supplementing limitations in the hosts translational machinery to ensure that phage particles are produced with high efficiency upon induction.

### Genome Architecture

Phages have a mosaic genome architecture (55, 56), in which the genome is organized into groups or clusters of genes. Mosaicism has been proposed to be driven by illegitimate recombination, but it is not clear from previous genomic analyses whether the mosaic boundaries are between clusters of related genes or between individual genes (32, 53, 57).

We identified the fourteen most commonly occurring and well-annotated genes within the prophage dataset (Table 1). The majority (66.9%; 10,542/15,765) of these predicted prophages had the integrase gene located near one of the ends of the prophage genome. We generated a genome data set where all the genomes were organized with the integrase at the same end of the genome, in some cases by reverse-complementing the genome to place the integrase gene consistently at the left end of every genome, and in other cases by recircularizing the genome *in silico* and introducing a break that placed the integrase at the left end.

The resulting alignment allowed us to compare genome architecture across all of the prophages, and revealed that gene order is conserved across phage genomes (Fig. 7a), suggesting that there is an advantageous order of clusters of related function within the genomes of prophages. To highlight the clusters, we used a modified Smith-Waterman approach, essentially ascribing a single letter to each of the functions and then aligning the genomes based on the order of the functions with a +1 score for a match and no penalty for a mismatch (Fig. 7b). This shows that the gene order across 1,537 phage genomes is highly conserved, reminiscent of the early—middle—late gene organization of many well characterized phages and eukaryotic viruses (58–60). We also visualized the presence of each gene at its location in the genome using a gene density plot (Fig. 8), which highlights the separation of the genes based on location in the genome. The combined results from these alignments strongly indicate that the mosaicism in phage genomes occurs at the gene-cluster level, and that phage genomes are not comprised of random hybrids of many genomes. Instead, recombination (illegitimate or otherwise) appears to be constrained to result in a highly conserved gene order where the early genes, including those encoding entry and DNA replication functions, precede those encoding packaging functions, which in turn precede tail formation and host lysis functions.

**Fig. 7.**
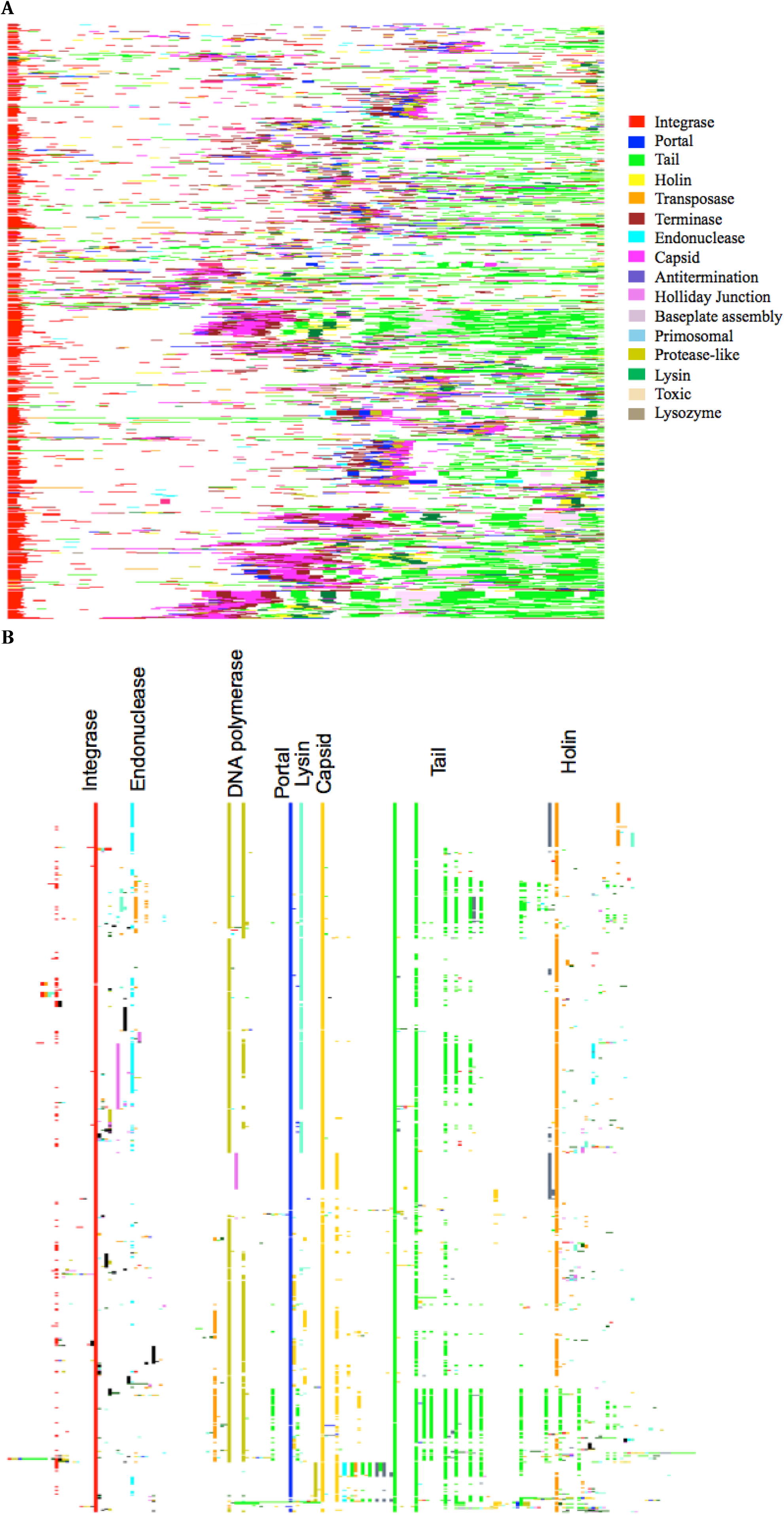
**(A)** Conserved prophage genome organization. Proteins in the phage genomes are colored by their annotation (integrase, portal, tail, holin, transposase, terminase, endonuclease, capsid, antitermination, Holliday junction resolvase, baseplate assembly protein, primosomal, protease-like, lysin, toxin-encoding genes, and lysozyme), and the phage genomes were normalized to the largest prophage. **(B)** Phage gene heatmap aligned using a modified Clustal alignment algorithm, as described in Materials and Methods.

**Fig. 8.**
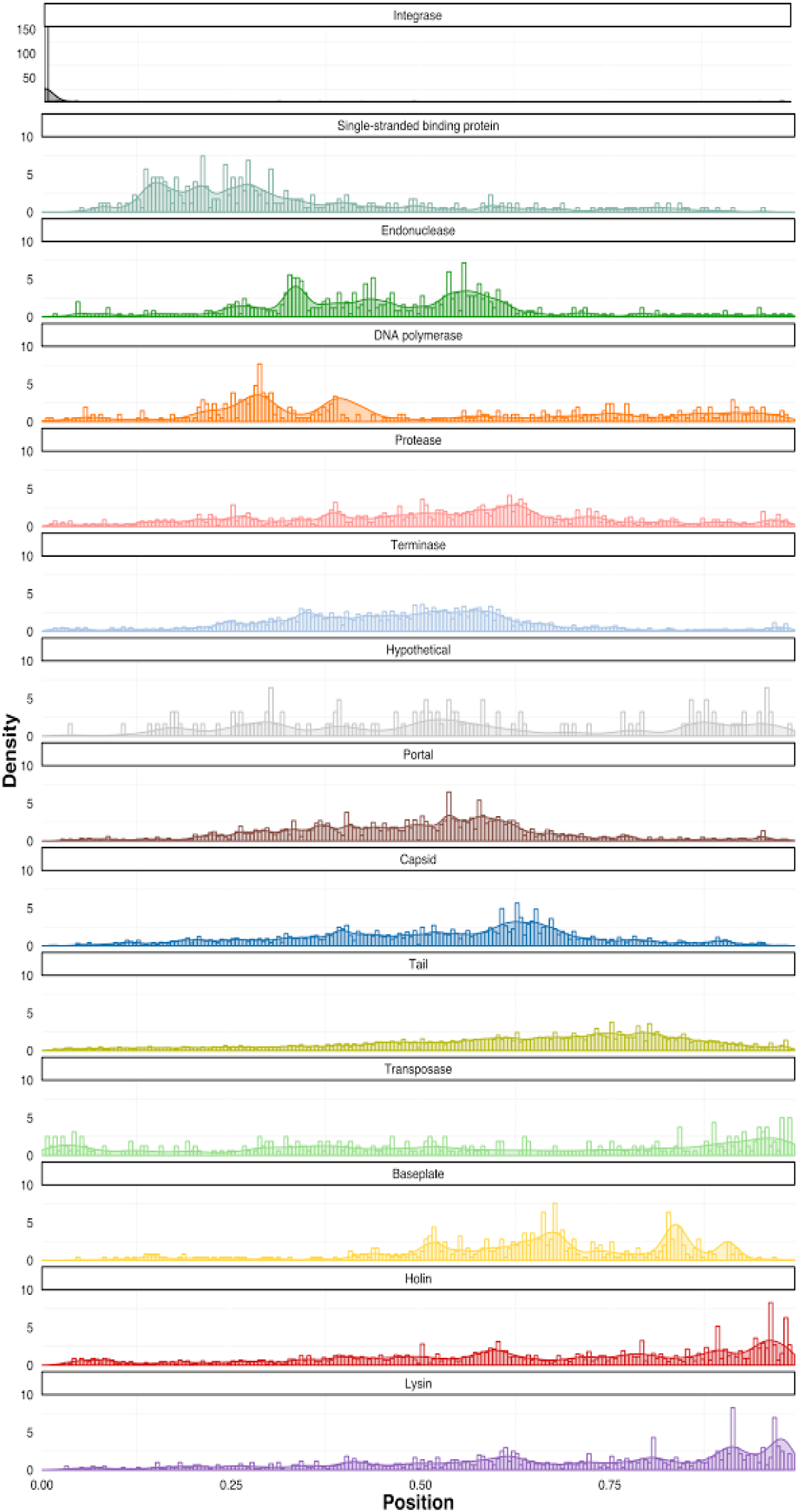
Gene density plots of the occurrence of phage marker genes along the genomes of 1,537 phage genomes. The genomes were rearranged as described in the text to place the integrase gene at the 5’ end of the phage genome, and then genes were displayed as ordered by the modal frequency of each gene.

The phage genome is therefore not a collection of genes that have been acquired at random by recombination, but rather analogous to a computer motherboard, where modules may be replaced provided they are slotted into the appropriate gene expression framework. Thus, a viable phage is more likely to be produced if genes are placed in the appropriate modules.

## Conclusions

Here we presented an analysis of over 11,000 bacterial genomes from which we identified 36,488 prophages. Many phages appear to be limited by initiation of translation by the host's machinery, and may increase translation rates by carrying their own tRNA genes, effectively increasing the availability of both tRNAs loaded with methionine and of peptide deformylase. We have also demonstrated that phages maintain a highly conserved gene order that suggests phage genome mosaicism is limited to clusters of conserved genes rather than individual genes.

